# Engineering “self-homing” circulating tumour cells as novel cancer theranostics

**DOI:** 10.1101/746685

**Authors:** Katie M Parkins, Veronica P Dubois, John J Kelly, Yuanxin Chen, Paula J Foster, John A Ronald

**Affiliations:** Robarts Research Institute, The University of Western Ontario, London, Ontario, Canada; The Department of Medical Biophysics, The University of Western Ontario, London, Ontario, Canada; Lawson Health Research Institute, London, Ontario, Canada

## Abstract

Purpose: New ways to target and treat metastatic disease are urgently needed. Tumor “self-homing” describes the recruitment of circulating tumor cells (CTCs) back to a previously excised primary tumor location, contributing to tumor recurrence, as well as their migration to established metastatic lesions. Recently, self-homing CTCs have been exploited as delivery vehicles for anti-cancer therapeutics in preclinical primary tumor models. However, the ability of CTCs to self-home and treat metastatic disease is largely unknown. Methods: Here, we employ molecular imaging to explore whether systemically-administered CTCs home to metastatic lesions and if CTCs armed with both a reporter gene and a cytotoxic prodrug gene therapy can be used to visualize and treat metastatic disease. Results: Bioluminescence imaging (BLI) performed over time revealed a remarkable ability of CTCs to home to primary and metastatic tumors throughout the body. Mice that received therapeutic CTCs had less BLI signal as well as less primary tumour burden than control mice. Preliminary data also showed self-homing therapeutic CTCs may be effective at treating disseminated breast cancer metastases. Conclusion: Using dual-luciferase BLI, this study demonstrates the noteworthy ability of experimental CTCs to home to disseminated breast cancer lesions. Moreover, by incorporating a prodrug gene therapy system into our self-homing CTCs, we show exciting progress towards effective and targeted delivery of gene-based therapeutics to treat both primary and metastatic lesions.

## Introduction

Cancer patient outcomes have significantly improved in the last few decades due to superior cancer imaging, surgical, and radiotherapy techniques, the recent application of ‘omics’ lesion profiling to guide therapy, and more efficacious drugs [1]. These advances now allow many patients with localized primary tumours or minimal metastatic disease at the time of diagnosis, or during recurrence, to be effectively managed. Yet despite these transformative advances, the ability to benefit patients with highly disseminated metastatic disease remains a significant challenge. Difficulties with controlling metastatic disease include, but are not limited to, the lack of tools to visualize lesions at an earlier stage when they may be more readily treated, insufficient systemic delivery of therapeutics to all lesions, and, most notably, extensive tumour heterogeneity both within and between lesions throughout the body [2, 3]. These substantial barriers highlight an unmet need for new technologies to effectively visualize and treat metastatic lesions, preferably so-called theranostic tools that have both diagnostic and therapeutic capabilities.

Cells are an attractive form of theranostic vector as they can be readily engineered *ex vivo* prior to transplantation with both molecular-genetic imaging reporter genes for noninvasive localization and therapeutic transgenes [4–7]. While some cell types have been shown to naturally home to lesions, such as stem cells and immune cells [8–14], one can also engineer cells with receptors targeting tumour-associated antigens to redirect *in vivo* cellular tropism. Recently, chimeric antigen receptor T cells (CAR-T cells) targeting the B cell antigen CD-19 became the first genetically-modified cell-based therapies to be approved for patients with relapsed or refractory B-cell precursor acute lymphoblastic leukemia and large B cell lymphoma [15–18]. While substantial efforts are now aimed at using CAR-T cells for the treatment of solid tumours, so far, their less than ideal therapeutic effectiveness has been attributed to insufficient tumour-homing and/or intratumoural immunological barriers [19]. Thus, the continued exploration of alternative cell types that can effectively home to metastatic solid tumours for use as novel theranostic vectors is warranted.

Paget’s “seed and soil hypothesis” describes the wide dissemination of “seeds”, or circulating tumour cells (CTCs), from a primary tumour and the formation of overt metastases selectively in “soils” that permit CTC survival and proliferation [20]. However, due to the non-permissive nature of tumour-free organs, metastasis has been shown to be an inefficient process in both experimental animal models and cancer patients [21–23]. The impedance of the formation of new metastases has been partly attributed to both vascular barriers that inhibit CTC extravasation from the blood as well as unfavorable survival conditions [24]. Conversely, shed CTCs have been shown to be highly capable of homing back to their tumour of origin, a concept termed tumour “self-seeding” that was first suggested and demonstrated by Kim and colleagues [25]. Self-seeding has been shown in animal models of human breast, colon and melanoma cancer, and is theorized to contribute to tumour recurrence following resection [25]. Unlike in tumour-free organs, tumour vasculature is often “leaky” due to a compromised vascular endothelium, and thus, more easily facilitates the extravasation of CTCs back into their original tumours [26]. Moreover, the primary tumour microenvironment is considered highly permissive soil for the continued survival and growth of recruited CTCs, leading to the expansion of highly metastatic clones that have a higher capacity to seed distant organs [25]. Similarly, metastatic lesions that have formed in distant organs are also considered fertile soil for additional “self-homing” CTCs to migrate to, survive, and expand within, which may contribute to accelerated metastatic disease progression [25].

In the last two decades, several groups have exploited self-homing CTCs as “self-targeted” delivery vehicles for *ex vivo* loaded anti-cancer therapeutic cargo [27–32]. Cargo has included oncolytic viruses such as the H-1 parvovirus and vesicular stomatitis virus (VSV), prodrug converting enzyme genes including herpes simplex virus thymidine kinase (HSV-TK) and cytosine deaminase (CD), transgenes that target the tumour microenvironment such as tumour necrosis factor (TNF), and the secretory version of TNF-related apoptosis-inducing ligand (S-TRAIL). Additionally, a few groups have co-engineered the therapeutic CTCs and/or their viral cargo with optical or positron emission tomography (PET) imaging reporter genes to enable the fate of the cells/cargo to be monitored with molecular-genetic imaging [28–30, 32]. Importantly, while the ability to target, visualize, and treat singular pre-established subcutaneous tumours as well as orthotopic or metastatic lesions in a singular organ (e.g., lungs [28] or brain [32]) has been demonstrated, to the best of our knowledge, the ability of self-homing CTCs to migrate into and be used to visualize and treat spontaneous multi-organ metastatic disease is largely unknown.

Here, we employed longitudinal molecular-genetic imaging to show that systemically-administered engineered CTCs efficiently home to both orthotopic and spontaneous metastatic breast cancer lesions. Further, we demonstrate that CTCs armed with both an imaging reporter gene and the gene for the prodrug converting enzyme cytosine deaminase-uracil phosphoribosyltransferase (CD:UPRT) can be used to effectively visualize and treat metastatic disease, resulting in distinctly increased survival times. Our preclinical study supports engineered CTCs as a novel self-targeting cellular theranostic platform for the visualization and treatment of distributed metastases - the most relevant lesions to patient outcome.

## Results

### Tracking of Self-Homing Cancer Cells in a Contralateral Orthotopic Tumour Model

Previous studies have shown that breast cancer cells from one mammary fat pad can home into a contralateral mammary fat pad (MFP) tumour [25]. Thus, we first started exploring the use of imaging to monitor tumour self-homing using this same experimental setup. We engineered the mouse breast cancer cell line (4T1) and its brain-seeking metastatic variant (4T1BR5) to express the orthogonal bioluminescence imaging (BLI) reporters *Renilla* luciferase (RLuc) and *Firefly* luciferase (FLuc), respectively. This allowed us to sensitively track both populations in the same animal over time. 4T1 cells were transduced with a lentiviral vector encoding both RLuc and ZsGreen and sorted to obtain 4T1-RLuc cells (Suppl. 1A). No significant change in ZsGreen expression over multiple passages was seen (Suppl. 1B) and there was a significant positive correlation shown between the number of 4T1-RLuc cells and RLuc/ZsGreen signal (R^2^ = 0.99, *p<0.001*; Suppl. 1C). FLuc-expressing 4T1BR5 (4T1BR5-FLuc) cells were engineered and characterized similarly in a previous study [33]. We next ensured a lack of cross-reactivity of the luciferase substrates. 4T1BR5-FLuc cells incubated with D-luciferin demonstrated significantly higher BLI signal than 4T1-RLuc cells, 4T1 parental cells, or equivalent volume of media, and 4T1-RLuc cells did not produce signal significantly different than 4T1 parental cells or media alone *(p<0.001*; Suppl.1D). Similarly, after the addition of h-Coelenterazine, 4T1-RLuc cells had significantly higher signal than 4T1BR5-FLuc cells, 4T1 parental cells, or equivalent volume of media and 4T1BR5-FLuc cells did not produce signal significantly different than 4T1 parental cells or media alone *(p<0.001*; Suppl. 1E). We next explored the migration of our engineered cells towards conditioned media from both cell lines using transwell migration assays. A significant increase in cell migration was seen for 4T1BR5-FLuc cells when conditioned media from 4T1-RLuc cells was used compared to conditioned media from 4T1BR5-FLuc cells or unconditioned media *(p<0.01*; Suppl. 1F). A significant increase in cell migration was also seen for 4T1-RLuc cells when conditioned media from 4T1-RLuc cells was used compared to unconditioned media *(p<0.01*; Suppl. 1F).

4T1-RLuc cells were then implanted into the right MFP of nude mice (n=5) and 4T1BR5-FLuc cells were implanted into the contralateral (left) MFP (Figure 1A). This allowed us to validate the lack of substrate cross-reactivity *in vivo* at early time points after cell injection (Days 0 and 1; Figures 1 and Suppl. 2) as well as the ability to evaluate whether either of the cell lines migrated into the contralateral MFP tumour (Figure 1). On Day 0 after cell injection, 4T1-RLuc cells only showed signal after administration with h-coelenterazine and on Day 1, 4T1BR5-FLuc cells only showed signal after administration of D-Luciferin (Suppl. 2). By day 7, 4T1BR5-FLuc cells did not appear to migrate as FLuc signal was not detected in the contralateral MFP (Figure 1D). In contrast, 4T1-RLuc cells could be detected in the contralateral MFP tumour, and RLuc signal was significantly higher in the contralateral compared to ipsilateral MFP (Figure 1C). The presence of both 4T1-RLuc and 4T1BR5-FLuc cells in the left MFP was confirmed histologically (Figure 1E, 1F), supporting our non-invasive imaging results and validating that the 4T1-RLuc cells left their initial site of implantation and homed to the contralateral MFP tumour.

**Figure 1:**
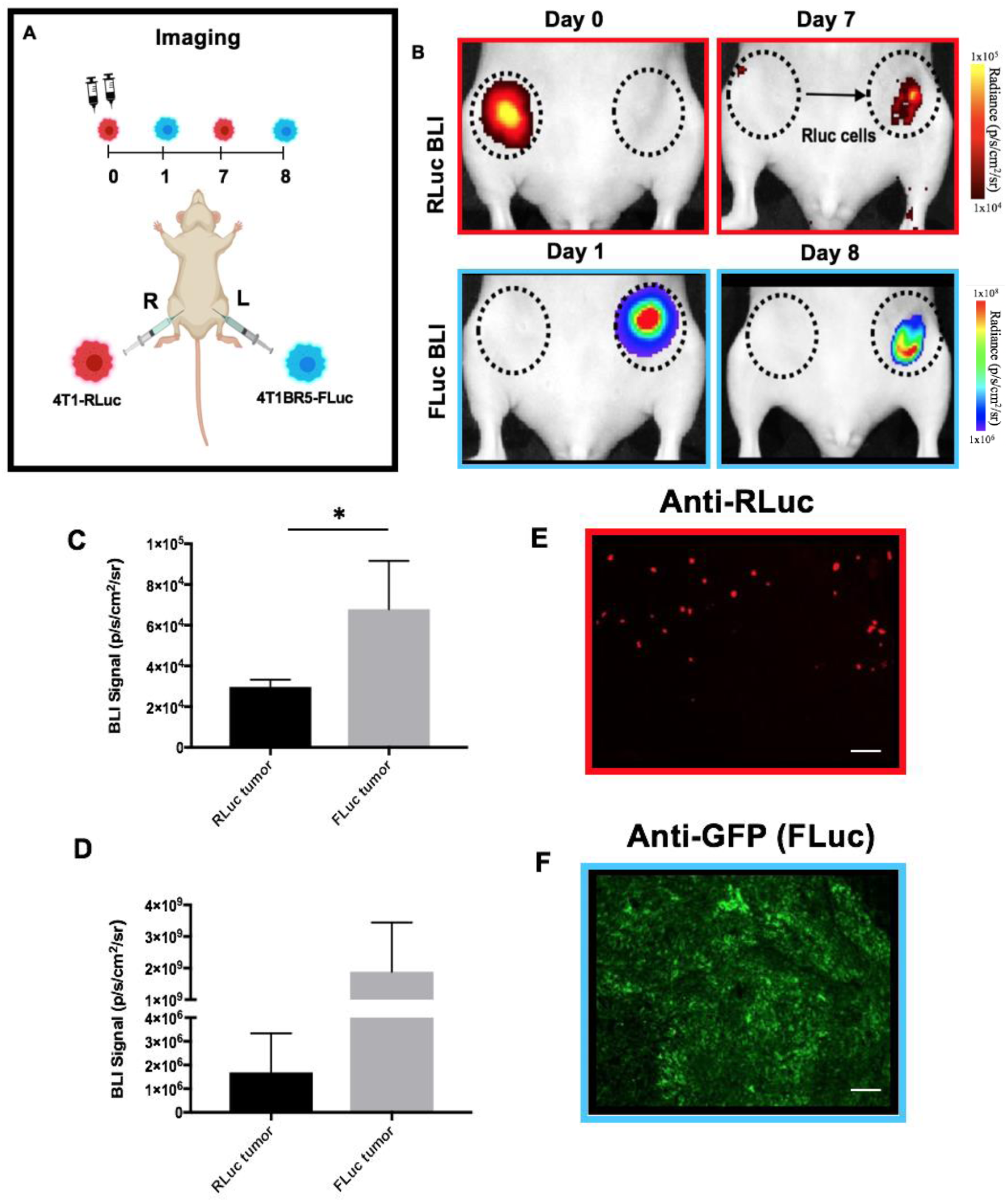
Experimental timeline for contralateral tumour self-homing model (n=5): On Day 0 after cell injection, 4T1-RLuc cells only showed signal after administration with h-coelenterazine and on Day 1, 4T1BR5-FLuc cells only showed signal after administration of D-Luciferin. By day 7, 4T1BR5-FLuc cells did not appear to migrate as FLuc signal was not detected in the contralateral MFP but 4T1-RLuc cells could be detected in the contralateral MFP tumour (B). RLuc signal on day 7 was significantly higher in the contralateral MFP compared to the ipsilateral MFP on day 8 (C/D). The presence of both 4T1-RLuc and 4T1BR5-FLuc cells in the left MFP was confirmed histologically (scale bars= 500 microns) (E/F).

### Monitoring Self-Targeted Therapy in a Contralateral Orthotopic Tumour Model

We next evaluated the ability of self-homing 4T1-RLuc cells expressing the therapeutic prodrug converting fusion enzyme cytosine deaminase-uracil phosphoribosyltransferase (CD:UPRT) to treat contralateral MFP tumours. CD:UPRT converts non-toxic 5’-fluorocytosine (5’FC) into the cytotoxic compound 5’fluoruridine monophosphate (5’FUMP) and was chosen as a suicide switch to eliminate the therapeutic cells as well as a way to kill adjacent non-engineered cancer cells via the bystander effect [34–37]. 4T1-RLuc and 4T1BR5-FLuc cells were transduced with a lentiviral vector co-expressing CD:UPRT (CD for brevity) and tdTomato (tdT), and sorted via tdT to obtain 4T1-RLuc/CD (4T1-CD) cells and 4T1BR5-FLuc/CD (4T1BR5-CD) cells (Figures 2A and Suppl. 3A). After 96 hours of incubation with 5’FC (5mM), CD expressing cells showed significantly less survival than cells without drug as well as significantly less survival than 4T1-RLuc and 4T1BR5-FLuc cells with or without drug (Figures 2B and Suppl. 3B). At all doses (0.005mM, 0.05mM, 0.5mM, 5mM), CD expressing cells show significantly less survival than cells without drug (Figures 2C and Suppl. 3C).

**Figure 2:**
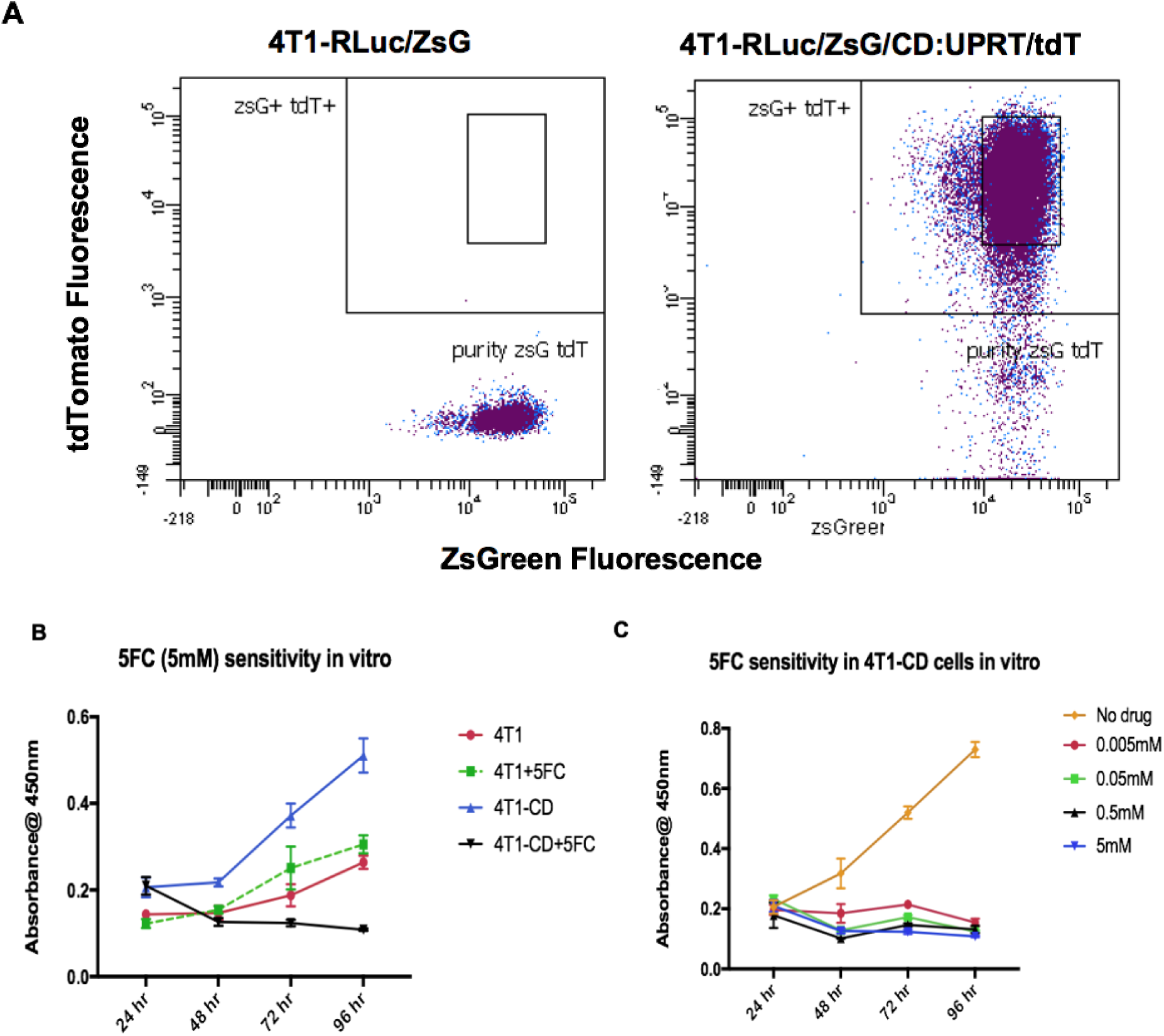
Therapeutic cell characterization: 4T1-RLuc cells were transduced with a lentiviral vector co-expressing the therapeutic prodrug converting fusion enzyme cytosine deaminase-uracil phosphoribosyltransferase (CD:UPRT) and tdTomato (tdT), and sorted via tdT to obtain 4T1-RLuc/CD cells (A). After 96 hours of incubation with 5’FC (5mM), CD expressing cells showed significantly less survival than cells without drug as well as significantly less survival than 4T1-RLuc cells with or without drug (B). At all doses, CD expressing cells show significantly less survival than cells without drug (C).

Next, we performed *in vivo* experiments whereby either 4T1-RLuc (n=4) or 4T1-CD (n=4) cells were implanted into the right MFP and 4T1BR5-FLuc into the contralateral MFP. All mice were treated with 5’FC daily from days 8 to 14 (Figure 3A). As visualized with RLuc BLI, on day 7 before treatment, both 4T1-CD and 4T1-RLuc cells can be seen in the contralateral MFP tumour and RLuc signal at this site was not significantly different between mouse cohorts (Figure 3B, 3C). At day 14 following treatment, FLuc BLI signal was observed in both contralateral and ipsilateral MFPs and signal was not significantly different between the two mouse groups in both MFPs (Figure 3E, 3F). At endpoint, areas of necrosis were evident in MFP tumours from both mouse cohorts using hematoxylin and eosin (H&E) staining (Figure 3G).

**Figure 3:**
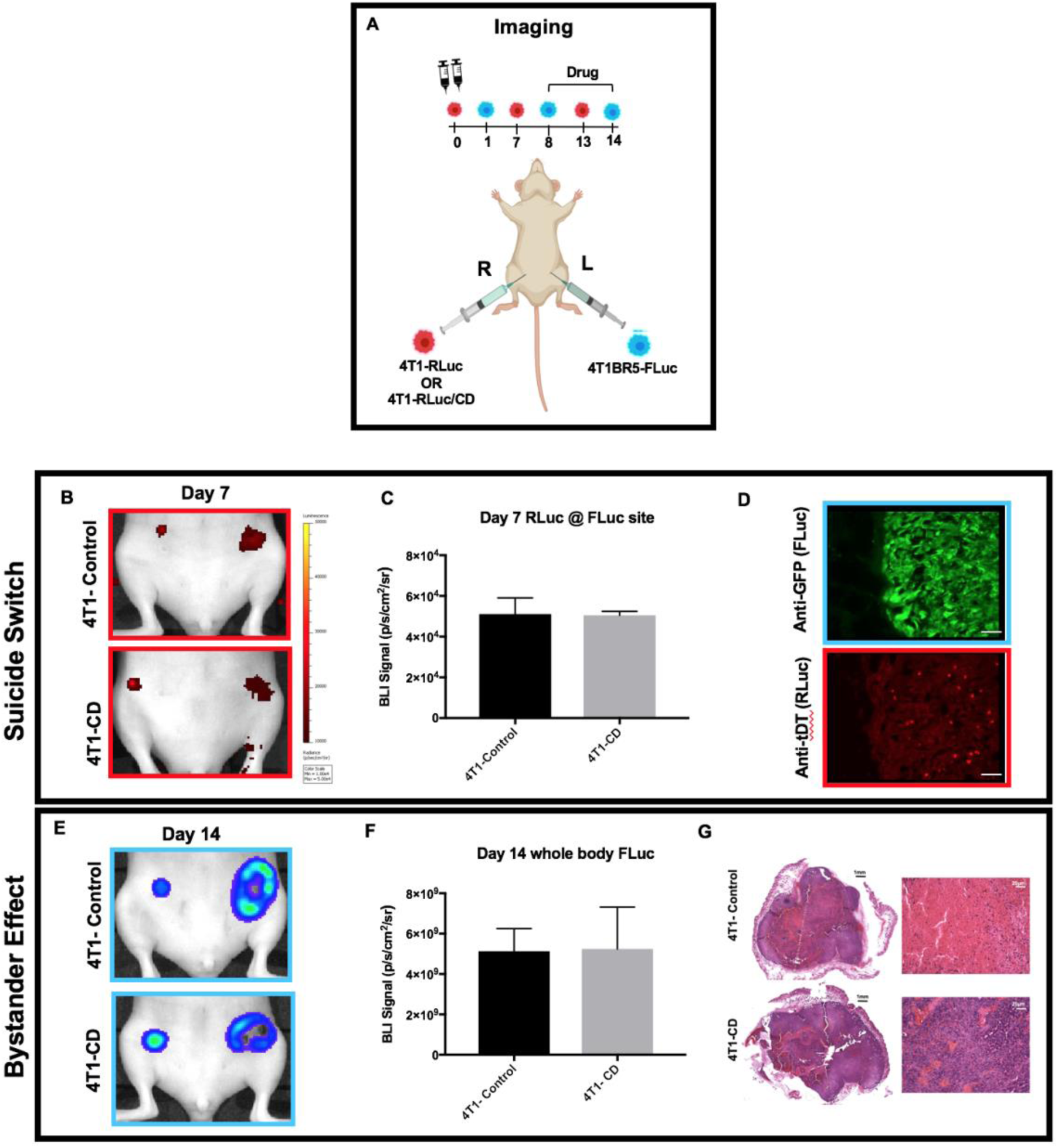
Experimental timeline for contralateral tumour treatment (n=8): All mice were treated with 5’FC daily from days 8 to 14. As visualized with RLuc BLI, on day 7 before treatment, both 4T1-RLuc/CD and 4T1-RLuc cells can be seen in the contralateral MFP tumour and RLuc signal at this site was not significantly different between mouse cohorts (B/C). tDT expressing therapeutic cells were visualized in the contralateral MFP using fluorescence microscopy at day 7 before drug administration (scale bars=200 microns). At day 14 following treatment, FLuc BLI signal was observed in both contralateral and ipsilateral MFPs and signal was not significantly different between the two mouse groups (E/F). At endpoint, areas of necrosis were evident in MFP tumours from both mouse cohorts using hematoxylin and eosin (H&E) staining (black scale bars= 1mm; white scale bars= 20 microns) (G).

### Intratumoural Injection of Therapeutic Cancer Cells Can Treat Orthotopic Tumours

We hypothesized the lack of therapeutic effect in the previous experiment may have been due to insufficient numbers of 4T1-CD cells migrating into the contralateral MFP tumour. Immunostaining of mice sacrificed on day 7 confirmed the presence of both 4T1BR5-FLuc and 4T1-CD cells in the left MFP, but a relatively low ratio of 4T1-CD to 4T1BR5-FLuc cells was noted (Figure 3D). To further test our hypothesis, we allowed the 4T1BR5-FLuc tumours to grow for 7 days prior to injecting 3×10^5^ 4T1-RLuc or 4T1-CD cells intratumourally and treated all mice with 5’FC daily from days 8 to 16 (Figure 4). 4T1 cells were visualized with RLuc BLI on days 7 and 15, and 4T1BR5 cells with FLuc BLI on days 0, 4, 8 and 16 (Figure 4A). Importantly, RLuc BLI of mice intratumourally injected with 4T1-CD cells was significantly higher than in mice that had the same cells home from the contralateral MFP the day prior to treatment initiation (Figure 4B, 4C). At day 15 after treatment, mice that received 4T1-CD cells intratumourally had a larger percent signal loss of RLuc signal compared to mice that received 4T1-RLuc cells, indicating the ability to mitigate therapeutic cancer cell growth via suicide switch activation (Figure 4B, 4D). Furthermore, by day 16, mice that received 4T1-CD cells had significantly less FLuc signal compared to mice that received 4T1-RLuc cells, indicating kill of adjacent cells via the bystander effect (Figure 4E, 4F). Measuring FLuc signal by BLI was complicated by the development of tumour ulcerations in both groups, which partially blocked signal. Large areas of necrosis were seen in histological sections of MFP tumours from both mouse cohorts (Figure 4G). Therefore, we also assessed treatment response by measuring MFP tumour volumes over time with calipers. Mice that received 4T1-CD cancer cells had significantly smaller tumour volumes than control mice at both days 14 and 17 (Figure 4H, 4I).

**Figure 4:**
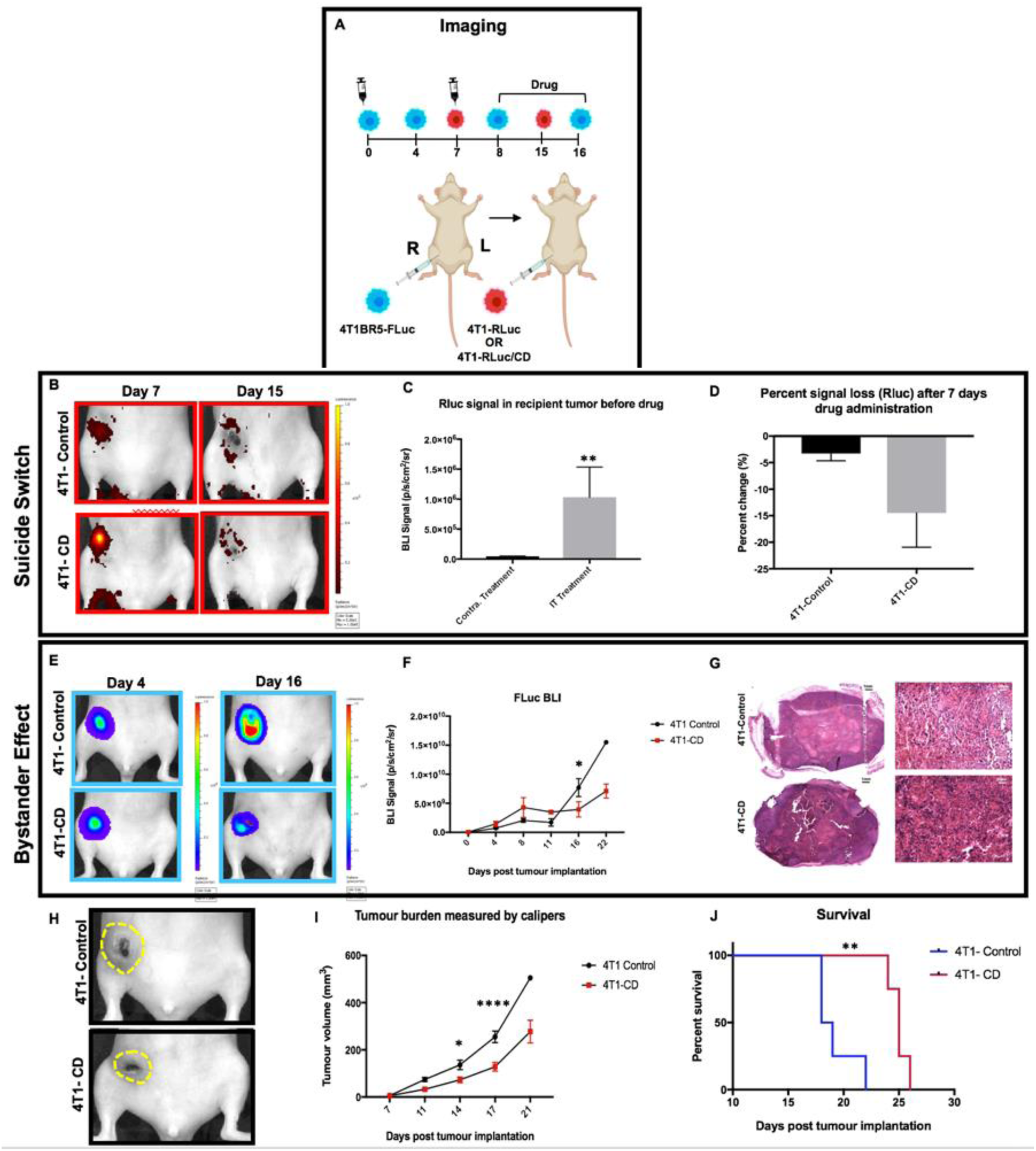
Experimental timeline for intratumoural injection of therapeutic cancer cells (n=8): All mice were treated with 5’FC daily from days 8 to 16. 4T1-Rluc and 4T1-RLuc/CD cells were visualized in the left MFP with BLI on days 7 and 15 (B) RLuc BLI of mice intratumourally injected with 4T1-RLuc/CD cells was significantly higher than in mice that had the same cells home from the contralateral MFP the day prior to treatment initiation (C). Mice that received 4T1-RLuc/CD cells intratumourally had a larger percent signal loss of RLuc signal compared to mice that received 4T1-RLuc cells, indicating the ability to mitigate therapeutic cancer cell growth via suicide switch activation (D). Furthermore, by day 16, mice that received 4T1-RLuc/CD cells had significantly less FLuc signal compared to mice that received 4T1-RLuc cells, indicating kill of adjacent cells via the bystander effect (E/F). At endpoint, areas of necrosis were evident in MFP tumours from both mouse cohorts using hematoxylin and eosin (H&E) staining (black scale bars= 1mm; white scale bars= 20 microns) (G). Treatment response was also assessed by measuring MFP tumour volumes over time with calipers. Mice that received 4T1-RLuc/CD cancer cells had significantly smaller tumour volumes than control mice at both days 14 and 17 (H/I). 5’FC treated mice that received 4T1-RLuc/CD cells showed significantly improved survival times compared to mice receiving 4T1-RLuc cells (J).

Of the 4 control mice, 3 had to be sacrificed prior to the day 21 due to predetermined endpoints, either the size of the tumour (> 2cm^3^) and/or the presence of excessive ulceration. As a result, 5’FC treated mice that received 4T1-RLuc/CD cells showed significantly improved survival times compared to mice receiving 4T1-RLuc cells (p<0.01; Figure 4J).

### Primary Tumours and Spontaneous Metastases can be Visualized with Systemically-Administered “Diagnostic” CTCs

We next assessed the ability of systemically-administered CTCs to home to primary tumours and spontaneous metastases. We implanted 4T1-RLuc cells into the right MFP of nude mice (n=5) and allowed tumours to grow for 7 days prior to injecting 4T1BR5-FLuc CTCs via an intracardiac injection under ultrasound guidance (Figure 5A). RLuc BLI was performed on days 0, 6, 13 and 19 to visualize cells in the right MFP and any spontaneous metastases and FLuc BLI was performed on days 7, 14 and 20 to visualize CTCs (Figure 5A, 5B). RLuc BLI showed the presence of metastases in 1 of 5 mice on day 6 prior to CTC injection. RLuc tumours were often found in the brain and/or hind limbs. FLuc BLI over time revealed the ability of FLuc-expressing CTCs to home to RLuc-expressing primary tumours and spontaneous metastases throughout the body (Figure 5B). Quantitative analysis of endpoint BLI images (day 19 and 20) revealed that the vast majority of metastases were composed of both 4T1-RLuc and 4T1BR5-FLuc cells (M=9.8±1.9), which was significantly higher than the number of metastases that were either 4T1-RLuc-positive only (M=0.6±0.4; *p<0.01*) or 4T1BR5-FLuc-positive only (M=0.2±0.2; *p<0.001*) (Figures 5C, 5D, Suppl. 4). The presence of both 4T1-RLuc and 4T1BR5-FLuc cells in numerous metastases was confirmed histologically (Figures 5E, Suppl. 4B), supporting our non-invasive imaging results.

**Figure 5:**
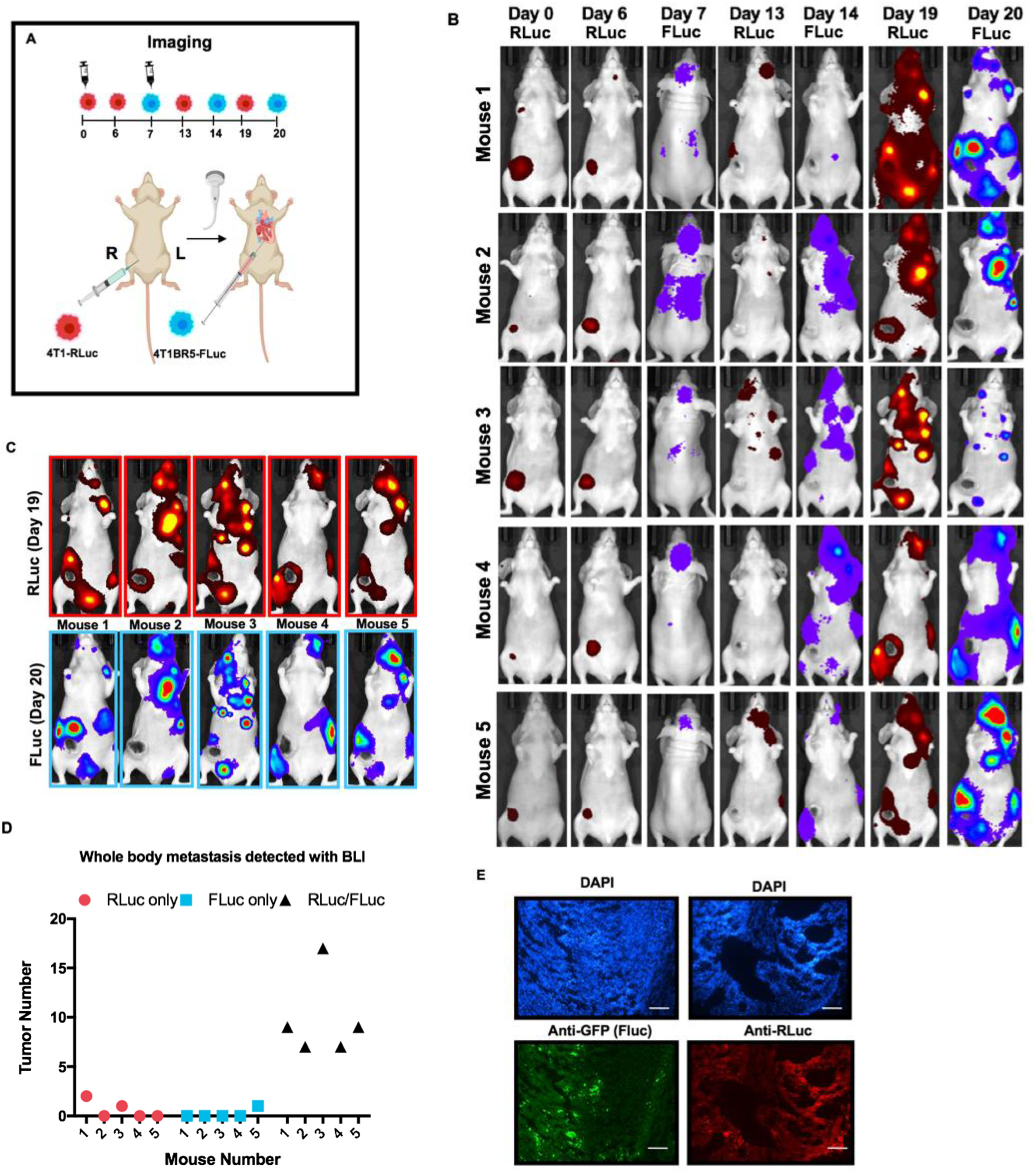
Experimental timeline for visualizing diagnostic CTCs (n=5): RLuc BLI was performed on days 0, 6, 13 and 19 to visualize cells in the right MFP and any spontaneous metastases and FLuc BLI was performed on days 7, 14 and 20 to visualize CTCs (B). FLuc-expressing CTCs efficiently homed to RLuc-expressing primary tumours and spontaneous metastases throughout the body (C). Quantitative analysis of endpoint BLI images (day 19 and 20) revealed that the vast majority of metastases were composed of both 4T1-RLuc and 4T1BR5-FLuc cells, which was significantly higher than the number of metastases that were either 4T1-RLuc-positive only or 4T1BR5-FLuc-positive only (p< 0.001) (C/D). The presence of both 4T1-RLuc and 4T1BR5-FLuc cells in metastases was confirmed histologically (scale bars= 500 microns) (E).

### Self-Homing “Theranostic” CTCs Efficiently Migrate into and Treat Primary Tumours and Spontaneous Metastases

Finally, we investigated whether theranostic CTCs could be systemically administered to treat primary and disseminated lesions. Again, 4T1-RLuc cells were implanted into the right MFP of nude mice and allowed 7 days of tumour growth prior to intracardiac injection of either 4T1BR5-CD cells (n=6) or 4T1BR5-FLuc cells (n=6). These mice were treated with 5’FC daily from days 10 to 20. We also included an additional cohort of mice (n=4) who received an MFP injection of 4T1-RLuc cells only and were not administered 5’FC (“4T1 Only mice” in Figure 6). In our first cohort (n=4), all mice had primary tumour growth similar to our data with our “diagnostic” CTCs and also developed sufficient metastases to assess therapeutic effects on metastases (Figure 7). In our second cohort, 12/12 mice had MFP tumours that did not develop as well as in previous cohorts at the time of secondary injection and thus, did not develop more than one or two metastases per mouse by endpoint. For these mice, we only assessed the effects of treatment on primary tumour growth (Figure 6). RLuc BLI was performed on days 0, 6, 13 and 19 to visualize cells in the right MFP and any spontaneous metastases and FLuc BLI was performed on days 7, 14 and 20 to visualize CTCs (Figure 6A). As visualized by RLuc BLI, by day 6 (prior to drug administration), mice receiving 4T1BR5-CD cells had MFP RLuc signal that was not significantly different than mice receiving 4T1BR5-FLuc cells or mice receiving 4T1 cells only (Figure 6B, 6C). However, by day 19, mice receiving 4T1 cells only, had significantly higher RLuc signal in the MFP compared to mice that received 4T1BR5-CD cells (Figure 6B, 6D). At endpoint, primary tumours were not palpable in mice that received 4T1BR5-CD expressing cells (Figure 6E, 6F). These data support the notion that systemically administered “theranostic” CTCs are capable of returning to and effectively treating the primary tumour. Of the 6 mice that received 4T1BR5-FLuc cells, 2 had to be sacrificed prior to endpoint due to both the size of the tumours and presence of ulceration. Overall, mice that received 4T1BR5-CD cells showed significantly improved survival compared to mice receiving 4T1BR5-FLuc cells and mice receiving 4T1 cells only (p<0.01; Figure 6G). In the 4 mice that developed notable metastases, mice receiving 4T1BR5-CD CTCs (n=2) and 4T1BR5-FLuc CTCs (n=2) had near equivalent FLuc signal on the day of CTC injection, but by day 14, mice receiving 4T1BR5-CD CTCs had visibly less FLuc signal (Figure 7B, 7C). Both mice that received 4T1BR5-FLuc cells had to be sacrificed prior to endpoint due to both the size of primary tumours and presence of ulceration, and thus, FLuc imaging on days 20 and 29 was only performed on mice that received 4T1BR5-CD cells. Importantly, mice in each cohort had similar RLuc signal on days 0 and 6 prior to CTC administration (Figure 7D, 7E), but by day 19, the single mouse that received 4T1BR5-FLuc cells and survived until day 19, had visibly more RLuc metastases than both mice that received 4T1BR5-CD cells (Figure 7D, 7E).

**Figure 6:**
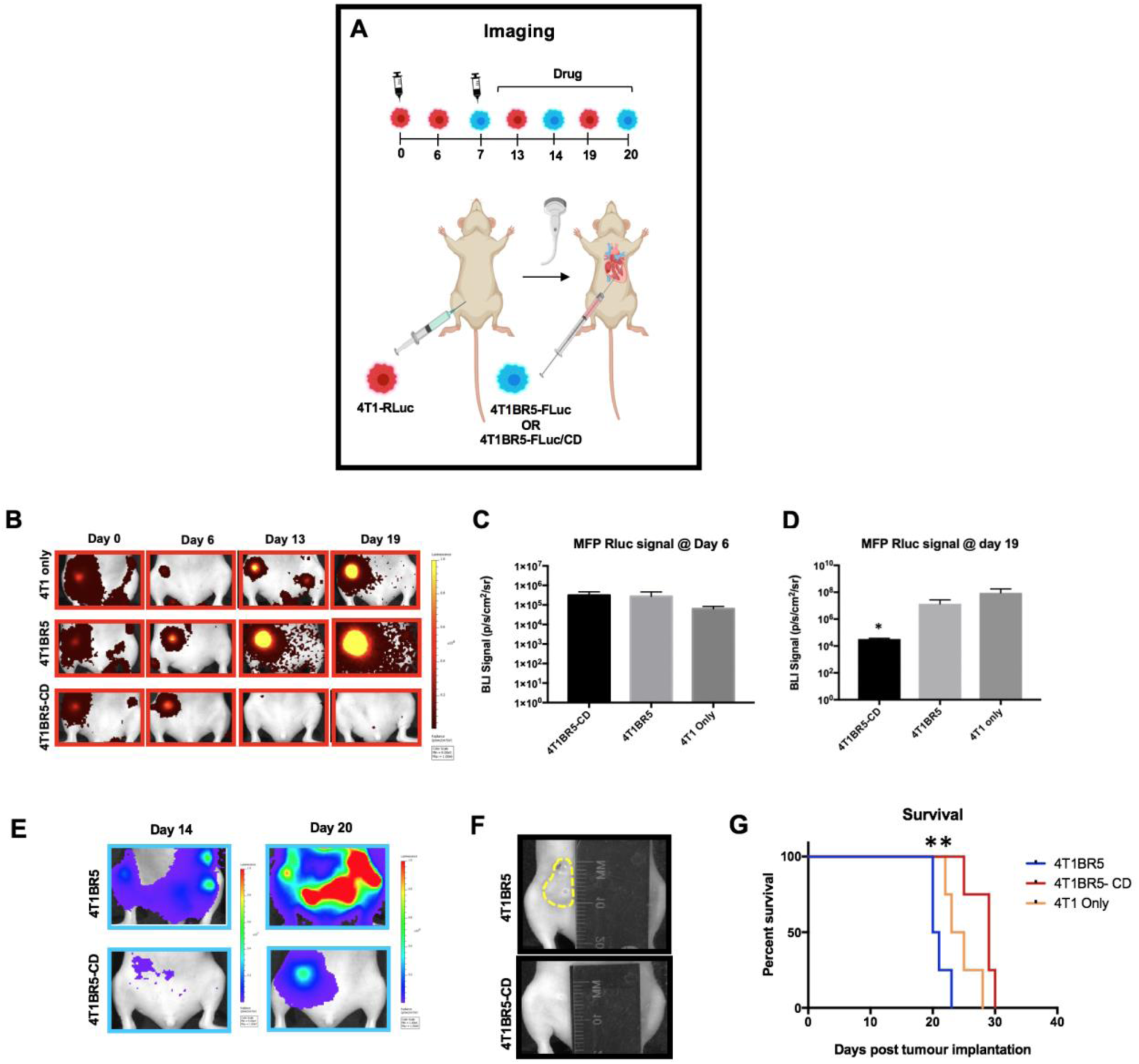
Experimental timeline for visualizing self-homing theranostic CTCs (n=16): 4T1BR5-CD and 4T1BR5 mice were treated with 5’FC daily from days 10 to 20. RLuc BLI was performed on days 0, 6, 13 and 19 to visualize cells in the right MFP and FLuc BLI was performed on days 7, 14 and 20 to visualize CTCs (B/E). By day 6 (prior to drug administration), mice receiving 4T1BR5-CD cells had MFP RLuc signal that was not significantly different than mice receiving 4T1BR5 cells or 4T1 cells only (C). However, by day 19, mice receiving only 4T1-RLuc cells had a significantly higher percent increase of RLuc signal in the MFP compared to mice that received 4T1BR5-CD cells (D). At endpoint, primary tumours were not palpable in mice that received 4T1BR5-CD cells (F). These data support the notion that systemically administered “theranostic” CTCs are capable of returning to and effectively treating the primary tumour (E). Mice that received 4T1BR5-CD cells showed significantly improved survival compared to mice receiving 4T1BR5 cells and mice receiving 4T1 cells only (G).

**Figure 7:**
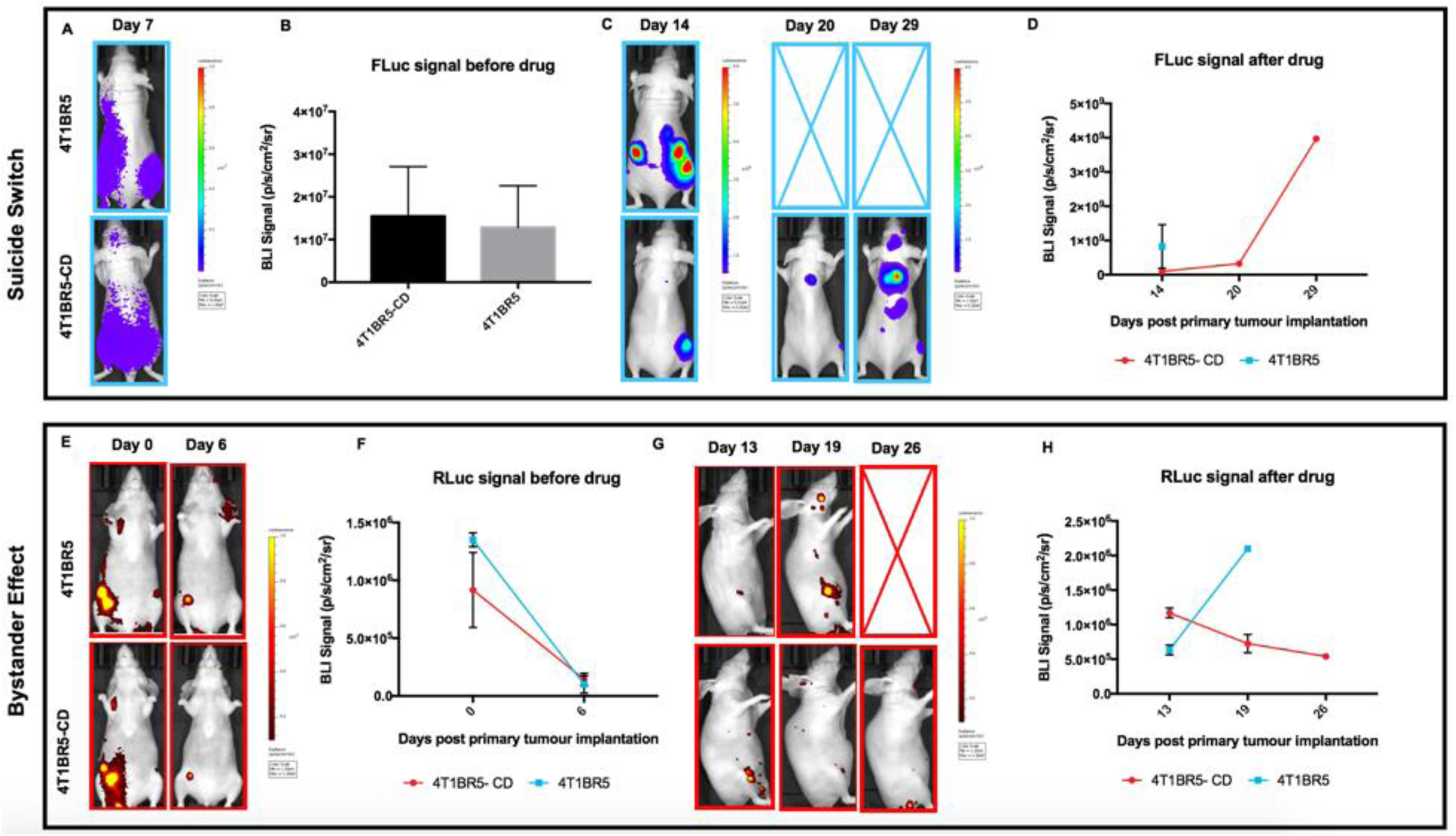
Treating metastases: On the day of intracardiac injection (day 7), mice receiving 4T1BR5-CD CTCs (n=2) and 4T1BR5 CTCs (n=2) had near equivalent FLuc signal (A/B), but by day 14, mice receiving 4T1BR5-CD CTCs had visibly less FLuc signal (C/D). Both mice that received 4T1BR5 cells had to be sacrificed prior to endpoint due to both the size of primary tumours and presence of ulceration, and thus, FLuc imaging on days 20 and 29 was only performed on mice that received 4T1BR5-CD cells. Importantly, mice in each cohort had similar RLuc signal on days 0 and 6 prior to CTC administration (E/F), but by day 19, the single mouse that received 4T1BR5 cells and survived until day 19, had visibly more RLuc metastases than both mice that received 4T1BR5-CD cells (G/H).

## Discussion

This study demonstrates that engineered “self-homing” CTCs co-expressing an imaging reporter and a therapeutic transgene can be used as a novel theranostic cellular vector to visualize and treat both primary tumours and disseminated spontaneous breast cancer metastases in mice. We first show using dual-luciferase BLI the remarkable ability of systemically-administered CTCs to preferentially home to pre-established spontaneous metastases in various organs throughout the body with minimal formation of new tumours. Leveraging on this highly preferential homing capability, we then show that CTCs co-expressing the prodrug converting fusion enzyme system CD:UPRT, to kill neighboring cells via the bystander effect as well as act as a suicide switch, was able to decrease tumour burden and extend survival.

Cancer, particularly in patients with metastatic disease, remains a leading cause of death in the world [38, 39]. Treatments that often work on localized disease are often not an option or fail in the patients with significant metastatic spread. Thus, the development of technologies for earlier detection and treatment of metastatic disease remains at the forefront of cancer research. Therapeutics that have natural tumour tropism or are designed to target lesions offer potential benefits of improved therapeutic effectiveness due to high intratumoural concentration of therapeutic payloads, as well as increased safety due to minimization of off-target cytotoxicity in normal tissues [4-19, 27-32]. The incorporation of imaging probes (e.g., radiolabels, iron oxides, etc.) into these targeted agents has been used to allow one to track their whole-body biodistribution to on-tumour and off-tumour sites [28–30, 32]. An agent that is very effective at naturally localizing to metastases with little off-target accumulation could theoretically also be used as an effective agent for the early diagnosis and treatment of metastatic disease.

In recent years, numerous classes of tumour-targeting agents have been developed including small molecules [40], antibodies [41–43], nanoparticles [44, 45], viral [46–48] and non-viral gene vectors [49–51], and cell-based vectors [4-19, 27-32]. Traditionally, the development of cell-based therapies has focused on the use of stem-cells including tissue-specific stem cells, pluripotent stem cells or mesenchymal stem cells [5,6,8–14]. These approaches offer many important benefits including their innate homing capabilities and natural anti-tumour effects; however, they suffer from limited passaging in culture resulting in roadblocks throughout the engineering process as well as limited therapeutic potential *in vivo*. Similarly, others have used immune cell-based vectors in efforts to target and treat a variety of tumours including tumour-infiltrating lymphocytes (TILs) [52, 53] and cytokine induced killer (CIK) cells [54, 55]. CIK cells are a group of immune effector cells with a mixed T- and natural killer (NK) cell phenotype. Thorne and colleagues successfully showed this effector cell population could be loaded with an oncolytic virus for targeted delivery and treatment of murine tumours [55]. Currently, one of the most employed cell-based tumour-targeting strategies is the engineering of T lymphocytes with chimeric antigen receptors (CARs) in efforts to selectively recognize and kill cancer cells expressing the B cell antigen CD19, while sparing healthy tissue [15–18]. This strategy has shown tremendous promise in the clinic in upwards of 70% of patients with otherwise treatment-refractory B cell leukemia and other B cell malignancies such as diffuse large B-cell lymphoma. However, in some cases, CAR-T cell therapy has shown no effect at all, and in many cases, patients experience potentially lethal side effects as a result of CAR-T cells not proliferating and persisting in the body over time, proliferating and activating excessively, or nonspecific homing to healthy organs [19, 56]. CAR-T cells have also shown minimal therapeutic effectiveness for solid tumours due to less than ideal homing capacity. Thus, there remains a need for novel cell-based delivery vehicles that are capable of homing to solid tumours.

In 2009, Kim and colleagues demonstrated the ability of cancer cells from an established lesion to enter the circulation and then return to this lesion to continue to survive and proliferate, a process they termed “self-seeding” [25]. The self-seeding capabilities of cancer cells was attributed to both the recruitment potential of the established tumour microenvironment as well as the seeding capabilities of cancer cells themselves. As a result, several groups have attempted to repurpose the self-homing properties of cancer cells to use them as “self-targeted” cell-based delivery vehicles for anti-cancer therapeutics [27–32]. In comparison to previously discussed cell-based vectors (i.e., stem cells and immune cells), cancer cells can be continuously grown *in vitro* enabling extensive cell engineering, may have superior homing capability to lesions, and may have prolonged survival and expansion once in tumours. Past studies have included using cancer cells as a vehicle to deliver oncolytic viruses [27, 28], engineering cancer cells to express suicide genes [29, 30] or apoptosis-inducing ligands [32] to transfer death signals to neighboring non-engineered tumour cells, and engineering cancer cells to express therapeutic agents that influence the tumour microenvironment [31], specifically angiogenesis. In this work, we exploit the innate self-homing properties of CTCs to investigate their potential as an efficient diagnostic probe and drug delivery vehicle for self-targeted therapy of primary and metastatic tumours. In our model, the primary tumour had a week to grow and spontaneously metastasize prior to the injection of experimental CTCs and thus, we assume it is the experimental CTCs homing to those pre-established sites and not the alternative. Our imaging data from early time points supports this theory such that some animals displayed spontaneous metastases prior to the injection of CTCs. Furthermore, our transwell migration assay data shows that conditioned media from our primary 4T1 cells, causes increased migration compared to media from our experimental CTC 4T1BR5 cell line. Previous work using the MDA-MB-231 breast cancer model showed that the cytokines IL-6 and IL-8 produced by the primary tumour acted as chemoattractants to efficiently recruit CTCs^25^. Future work will look to investigate whether the production of these cytokines is enhanced in 4T1 conditioned media and whether they may contribute to the self-homing efficiency seen in this model.

Based on our initial imaging results, we engineered our experimental CTCs to express a suicide gene and used noninvasive imaging to monitor the effects of self-targeted therapy in contralateral, intratumoural, and metastatic models. In a prodrug converting enzyme suicide gene therapy system, cancer cells are transfected with a gene that can express an enzyme with the ability to convert a nontoxic prodrug into an active chemotherapeutic [37]. By incorporating a suicide switch, we can visualize the killing of engineered therapeutic cancer cells as well as a bystander effect whereby adjacent non-engineered cancer cells are killed. While there are numerous enzyme/prodrug systems available, we chose to use the CD:UPRT fusion gene, due to the ability of 5’FU (the active form) to interrogate neighbouring cells independently of gap junctions, as well as readily diffuse the blood brain barrier [34–36]. As a result, the CD/5’FC system has shown to have a more potent bystander effect than the commonly used HSV-TK/GCV system [57–60]. Furthermore, in comparison to the traditional CD/5’FC system, we used the CD:UPRT fusion gene, as UPRT can further convert 5’FU into 5’FUMP, which has shown to have significantly enhanced cancer killing efficiency in prostate, ovarian and breast cancer subtypes [61, 62]. In the current study, we were able to visualize therapeutic CTCs that had successfully migrated from the original to the contralateral MFP but did not observe a significant therapeutic effect. However, we did observe a decrease in tumour burden when our therapeutic cells were administered intratumourally (3×10^5^ cells). These findings suggest the ratio of therapeutic cells to cancer cells may be crucial in generating a significant therapeutic effect *in vivo*. By using dual BLI, we also show effective homing and treatment of primary tumours when therapeutic CTCs were administered systemically and as a result, demonstrate extended survival in mice treated with 5’FC. Our preliminary data suggests systemically administered CTCs may also be capable of homing to and treating disseminated breast cancer metastases. Future work will look to validate these findings in additional mouse cohorts.

While our findings suggest CTCs have potential as highly-efficient carriers of therapeutic cargo to primary and metastatic tumour sites, our approach has some limitations to consider. Most importantly, while CD:UPRT expression was able to kill off many of the engineered 4T1BR5 cells, some of these cells were capable of avoiding cell death and generated new tumours. Additionally, while we demonstrate the ability to kill off neighbouring non-engineered cancer cells through a bystander effect, some mice show residual spontaneous metastases following treatment. Future work will explore incorporating more than one therapeutic gene into the engineered cells. For example, a CD/HSV-TK system, would allow the administration of two different pro-drugs, creating a higher likelihood of targeting and treating engineered CTCs while possibly also enhancing the bystander effect. Additionally, timing of CTC self-homing to pre-established tumours should be further explored. Our data suggests that three days may not be the optimal window for systemically administered CTCs to efficiently home to established metastases. If we administer the pro-drug too early, we may lose some of our therapeutic CTCs to self-induced toxicity prior to receiving any therapeutic effects on neighbouring non-engineered cancer cells. Furthermore, a larger window would allow time for therapeutic cells to proliferate and expand the therapeutic population once they have reached their tumour target.

In conclusion, our work provides evidence that CTCs are a novel theranostic vector platform for the visualization and treatment of pre-established tumour sites throughout the body. Overall, while further refinement is needed, this unorthodox strategy may have tremendous long term translational potential as a highly effective theranostic platform, specifically in patient populations presenting with metastatic disease at initial diagnosis, and those at high risk of cancer recurrence or metastatic relapse.

## Materials and methods

### Cell Engineering

The 4T1BR5 cells were a kind gift from Dr. Patricia Steeg’s lab and engineered to stably co-express red-shifted *Luciola italica* luciferase (FLuc) and green fluorescent protein (GFP) using a commercial lentiviral vector (RediFect Red-FLuc-GFP; PerkinElmer, USA). Cells were transduced and FACS sorted based on GFP expression using a FACSAria III flow cytometric cell sorter (BD Biosciences). The parental 4T1 cells were also received from Dr. Patricia Steeg’s lab and engineered to stably co-express *Renilla* luciferase-8 (RLuc) and ZsGreen (ZsG) using a virus made in house. Cells were transduced and sorted based on ZsG expression using FACS. The resultant 4T1BR5-FLuc/GFP (4T1BR5-Fluc) and 4T1-RLuc/ZsG (4T1-Rluc) cells were maintained in DMEM containing 10% FBS and 1% antibiotics, at 37°C and 5% CO_2_. The 4T1BR5-FLuc and 4T1-RLuc cells were then engineered a second time to stably express cytosine deaminase-uracil phosphoribosyl transferase (CD-UPRT) and tdTomato (tdT). Both cell lines were transduced and FACS sorted based on tdT expression. Cells were washed three times with Hanks balanced salt solution (HBSS) and collected for *in vitro* evaluation or injection into animals.

### In Vitro Studies

#### Cell line characterization

All *in vitro* results are from three independent experiments with three replicates of each condition. To evaluate the relationship between cell number and BLI signal, 1×10^4^, 5×10^4^, 1×10^5^, 1.5×10^5^, and 5×10^5^ 4T1BR5-FLuc or 4T1-RLuc cells were seeded in each well of 24-well plates. We acquired fluorescent images of each plate. We then added 10 μL of D-luciferin (30 mg/mL; Syd Labs, Inc., MA, USA) or 10 μL of h-Coelenterazine (150μg/mL; NanoLight Technology, Prolume, AZ, USA) to the growth medium in each well and BLI images were collected for up to 35 minutes. All images were acquired using a hybrid optical/X-ray scanner (IVIS Lumina XRMS In Vivo Imaging System, PerkinElmer). Signal was measured with region-of-interest (ROI) analysis using LivingImage Software (Perkin Elmer). An ROI was drawn around each well to measure the radiant efficiency (p/s/cm^2^/sr/uW/cm^2^) for fluorescence images and average radiance (p/s/mm^2^/sr) for bioluminescence images. The mean signal across replicates was determined for each independent experiment.

#### Cross reactivity

To assess *in vitro* cross reactivity, we seeded two identical 24-well plates with 1 x 10^5^ 4T1-RLuc, 4T1BR5-Fluc, 4T1 naïve cells, and equivalent volume of media. We added 10uL of d-Luciferin to each well in plate 1 and 10uL of h-Coelenterazine to each well in plate 2. Images were acquired for up to 35 minutes and an ROI was drawn around each well to measure the average radiance (p/s/mm^2^/sr). The mean signal across replicates was determined for each independent experiment.

#### Transwell Migration Assay

A FluoroBlok™ Multiwell Insert System was used with an 8um porous polyethylene terephthalate membrane (Corning, Corning NY, USA). We seeded 5×10^4^ cells (4T1-RLuc or 4T1BR5-Fluc) in 75cm^2^ flasks. At 48 hours post seeding, 650uL of new or conditioned DMEM was collected and used for the bottom chamber of the transwell plate. We then seeded 2.5×10^4^ cells (4T1-RLuc or 4T1BR5-Fluc) in the upper chamber of the transwell insert in 100ul of new DMEM. After 24 hours, the membranes were fixed in ethanol for 5 minutes, washed with PBS, and stained with Hoechst 33342 (10ug/ml in water) for 5 minutes. Membranes were cut out with a scalpel and mounted in 90% glycerol onto slides. Three random images were taken of the lower side of each membrane using an Invitrogen EVOS FL Auto Cell Imaging System and the mean fluorescence signal was calculated.

#### CD:UPRT Functionality Testing

To assess the functionality of the CD-UPRT gene *in vitro*, Vybrant MTT assays were used. 2×10^4^ 4T1-RLuc or 4T1-CD cells were seeded in each well of 96-well plates and incubated in either the desired concentration of 5’FC (diluted in DMEM) or incubated in DMEM alone. Ten microliters of MTT solution was added to each well and absorbance at 450nm was measured using a microplate spectrophotometer (Fluoroskan Ascent FL, ThermoLabSystems) at 24, 48, 72 and 96 hours. This experiment was repeated for 4T1BR5-FLuc and 4T1BR5-FLuc/CD cells.

### In Vivo Studies

Animals were cared for in accordance with the standards of the Canadian Council on Animal Care, and under an approved protocol of the University of Western Ontario’s Council on Animal Care (2015-0558). Six to eight-week-old female nu/nu mice were obtained from Charles River Laboratories (Willington, MA, USA).

#### Contralateral tumour model

Mice received a lower right mammary fat pad (MFP) injection of 300,000 4T1-Rluc or 4T1-CD cells and a lower left MFP injection of 300,000 4T1BR5-Fluc cells on day 0 (Figure 4.1A; n=5). RLuc BLI was performed on days 0 and 7 and FLuc BLI performed on days 1 and 8. Additional BLI was performed for experiments with CD expressing cells on days 15 (RLuc) and 16 (FLuc). For experiments with CD expressing cells, mice receiving 4T1-Rluc and 4T1-CD cells both received intraperitoneal injections of 5’FC (250mg/kg/day) on days 7 to 14 (Figure 4.3A; n=8).

#### Intratumoural model

Mice received a lower right mammary fat pad (MFP) injection of 300,000 4T1BR5-Fluc cells on day 0 and an intratumoural injection of 300,000 4T1-Rluc or 4T1-CD cells on day 7 (Figure 4.4A; n=8). FLuc BLI was performed on days 0, 4, 8 and 16 and RLuc BLI performed on days 7 and 15. Mice receiving 4T1BR5-Fluc and 4T1BR5-CD cells both received intraperitoneal injections of 5’FC (250mg/kg/day) on days 8 to 16.

#### Metastatic tumour model

Mice received a lower right MFP injection of 300,000 4T1-Rluc cells. MFP tumours grew for seven days prior to all mice receiving an intracardiac injection of 2×10^4^ 4T1BR5-FLuc or 4T1BR5-CD cells in 0.1mL of HBSS (Figure 4.5A; n=5). Injections were performed under image guidance using a Vevo 2100 ultrasound system (VisualSonics Inc.). RLuc BLI was performed on days 0, 6, 13 and 19. FLuc BLI was performed on days 7, 14 and 20. For experiments with CD expressing cells, mice receiving 4T1BR5-FLuc and 4T1BR5-CD cells both received intraperitoneal injections of 5’FC (250mg/kg/day) on days 10 to 20 (Figure 4.6A; n=12).

#### BLI Procedure

BLI was performed using a hybrid optical/X-ray scanner (IVIS Lumina XRMS In Vivo Imaging System, PerkinElmer). Mice were anesthetized with isofluorane (2% in 100% oxygen) using a nose cone attached to an activated carbon charcoal filter for passive scavenging. For RLuc BLI, anesthetized mice received a 20 μL intravenous injection of h-Coelenterazine (150μg/mL) and BLI images were captured for up to 30 minutes. For FLuc BLI, anesthetized mice received a 100 μL intraperitoneal injection of d-Luciferin (30 mg/mL) and BLI images were captured for up to 35 minutes.

#### Image Analysis

BLI signal was measured with region-of-interest (ROI) analysis using LivingImage Software (Perkin Elmer). ROIs were drawn throughout the mouse body of RLuc and FLuc image sets for each mouse.

#### Histology

At endpoint, mice were sacrificed by isoflurane overdose and perfused with 4% paraformaldehyde via the left ventricle. Tissues were removed and cryopreserved in ascending concentrations of sucrose (10, 20, and 30% w/v) for 24 hours each, then immersed in optimal cutting temperature (OCT) compound, and frozen using liquid nitrogen. Contiguous 10-μm frozen sections were collected and select sections were stained with hematoxylin and eosin (H&E), DAPI, Anti-GFP, Anti-Rluc. Stained sections were imaged using an Invitrogen EVOS FL Auto Cell Imaging System.

#### Statistics

All statistics were calculated using GraphPad Prism 7 Software. Data were expressed as mean ± SEM for *in vitro* and *in vivo* studies and analyzed by Student’s t test when comparing two groups. Survival times of mouse groups were analyzed using a log-rank test. Differences were considered statistically significant at *p < 0.05, **p < 0.01, ***p < 0.001, and ****p < 0.0001.

