## Supplemental Figures for "Engineering “self-homing” circulating tumour cells as novel cancer theranostics"

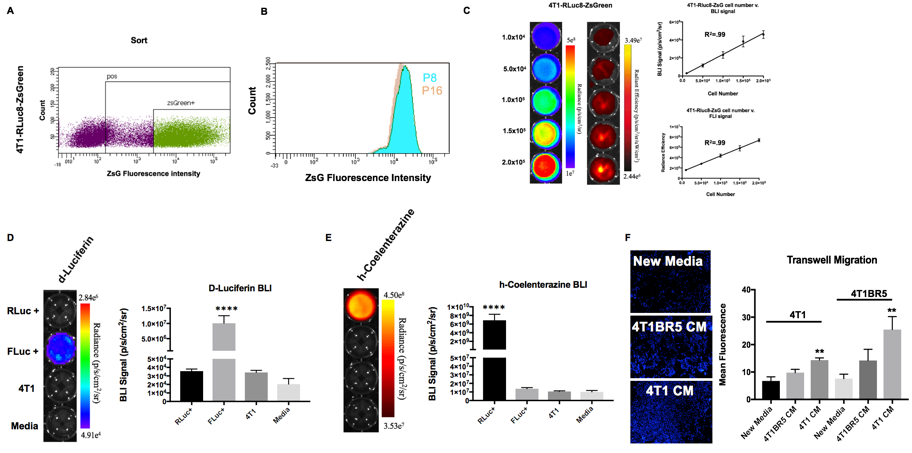

**Supplementary Figure 1: 4T1 cells were transduced with a lentiviral vector encoding both RLuc and ZsGreen and sorted to obtain 4T1-RLuc cells (Suppl. Fig. 1A). No significant change in ZsGreen expression over multiple passages was seen (B). There was a significant positive correlation shown between the number of 4T1-RLuc cells and RLuc/ZsGreen signal (C). 4T1BR5-FLuc cells incubated with D-luciferin demonstrated significantly higher BLI signal than 4T1-RLuc cells, 4T1 parental cells, or equivalent volume of media, and 4T1-RLuc cells did not produce signal significantly different than 4T1 parental cells or media alone *(*D). Similarly, after the addition of h-coelenterazine, 4T1-RLuc cells had significantly higher signal than 4T1BR5-FLuc cells, 4T1 parental cells, or equivalent volume of media and 4T1BR5-FLuc cells did not produce signal significantly different than 4T1 parental cells or media alone *(*E*)*. A significant increase in cell migration was seen for 4T1BR5 cells when conditioned media from 4T1 cells was used compared to conditioned media from 4T1BR5 cells or unconditioned media *(*F*).* A significant increase in cell migration was also seen for 4T1 cells when conditioned media from 4T1 cells was used compared to unconditioned media *(*F*).***

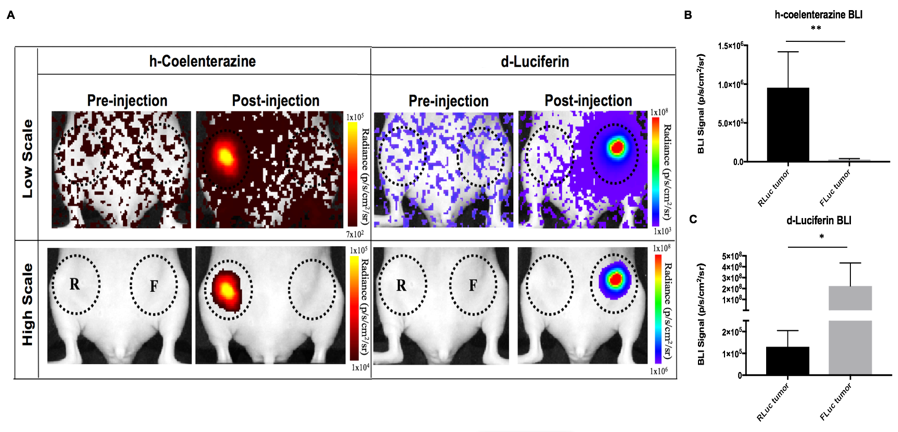

**Supplementary Figure 2: In vivo cross reactivity: 4T1-RLuc cells were implanted into the right MFP of nude mice (n=5) and 4T1BR5-FLuc cells were implanted into the contralateral (left) MFP (A). This allowed us to validate the lack of substrate cross-reactivity *in vivo* at early time points after cell injection. On Day 0, 4T1-RLuc cells only showed signal after administration with h-coelenterazine and signal in the right MFP was significantly higher than the left (B). Similarly, on Day 1, 4T1BR5-FLuc cells only showed signal after administration of d-Luciferin and signal in the left MFP was significantly higher than in the right MFP (C).**

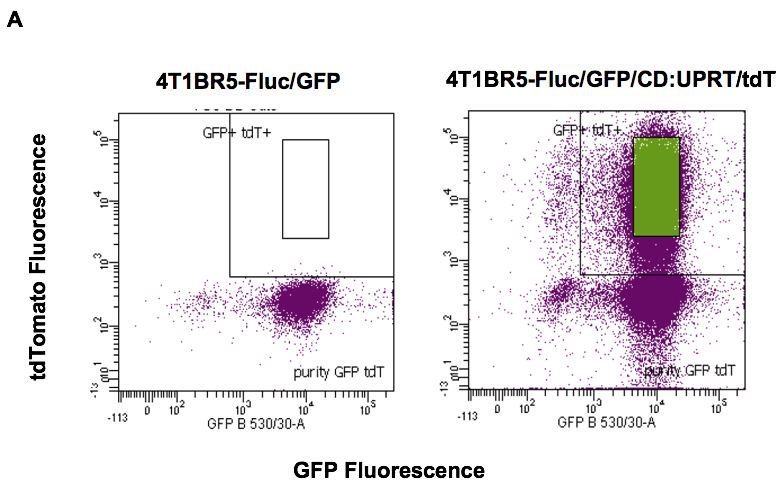

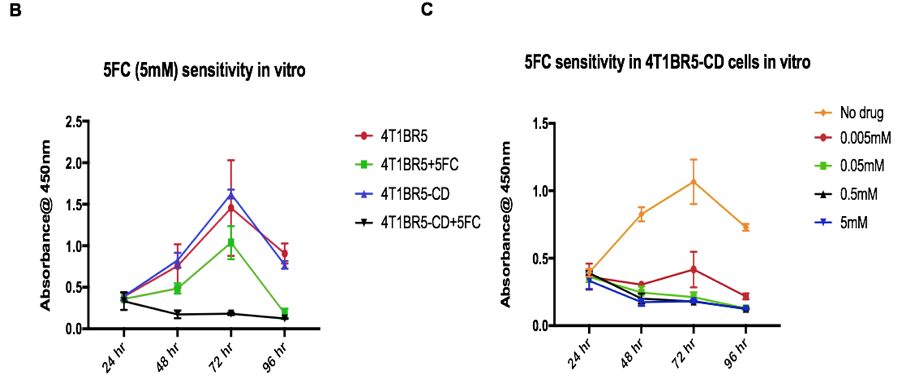

**Supplementary Figure 3: 4T1BR5-FLuc cells were transduced with a lentiviral vector co-expressing the therapeutic prodrug converting fusion enzyme cytosine deaminase-uracil phosphoribosyltransferase (CD:UPRT) and tdTomato (tdT), and sorted via tdT to obtain 4T1BR5-FLuc/CD cells (A). After 96 hours of incubation with 5’FC (5mM), CD expressing cells showed significantly less survival than cells without drug as well as significantly less survival than 4T1BR5-FLuc with or without drug (B). At all doses, CD expressing cells show significantly less survival than cells without drug (C).**

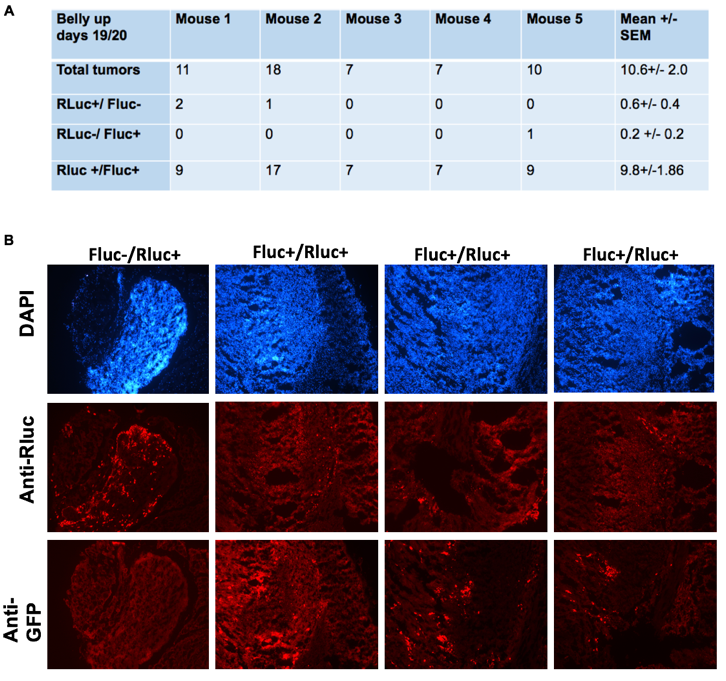

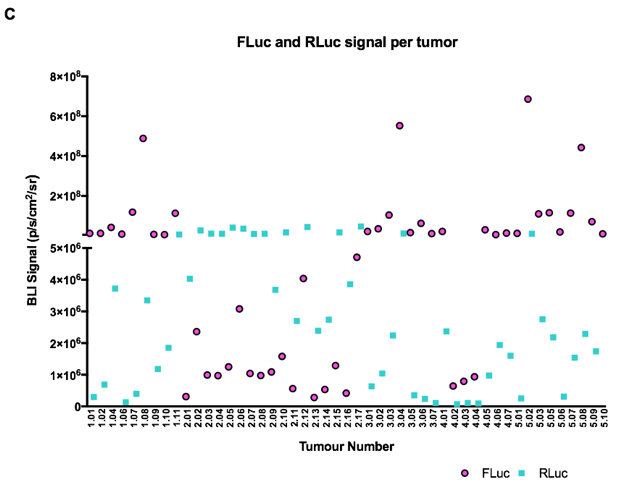

**Supplementary Figure 4: At endpoint, the number of metastases that were composed of both 4T1-RLuc and 4T1BR5-FLuc cells was significantly higher than the number of metastases that were either 4T1-RLuc-positive only or 4T1BR5-FLuc-positive only (A). The presence of both 4T1-RLuc and 4T1BR5-FLuc cells in numerous metastases was confirmed histologically (B). Using dual-BLI, we detected some whole-body metastases that had stronger FLuc signal than RLuc signal as well as metastases that had stronger RLuc signal than FLuc signal (X-axis values= mouse number followed by tumour number) (C).**
